# noSpliceVelo infers gene expression dynamics without separating unspliced and spliced transcripts

**DOI:** 10.1101/2024.08.08.607261

**Authors:** Tarun Mahajan, Sergei Maslov

**Affiliations:** Department of Bioengineering, University of Illinois Urbana-Champaign, Urbana, 61801, IL, USA; Center for Artificial Intelligence and Modeling, Carl R. Woese Institute for Genomic Biology, University of Illinois Urbana-Champaign, Urbana, 61801, IL, USA; Department of Physics, University of Illinois Urbana-Champaign, Urbana, 61801, IL, USA

**Keywords:** RNA velocity, Bursty gene expression, Cell state transitions, Burst size, Burst frequency

## Abstract

Modern single-cell transcriptomics has revolutionized biological research, but because of its destructive nature, it provides only static snapshots. Computational approaches that infer RNA velocity from the ratio of unspliced to spliced mRNA levels can be used to predict how gene expression changes over time. However, information about unspliced and spliced transcripts is not always available and may change on a timescale too short to accurately infer transitions between cellular states. Here we present noSpliceVelo, a novel technique for reconstructing RNA velocity without relying on unspliced and spliced transcripts. Instead, it exploits the temporal relationship between the variance and mean of bursty gene expression using a well-established biophysical model. When evaluated on datasets describing mouse pancreatic endocrinogenesis, mouse and human erythroid maturation, and neuronal stimulation in mouse embryonic cortex, noSpliceVelo performed comparably or better than scVelo, a splicing-based approach. In addition, noSpliceVelo inferred key biophysical parameters of gene regulation, specifically burst size and frequency, potentially distinguishing between transcriptional and epigenetic regulation.

## 1 Introduction

Single-cell transcriptomics has dramatically transformed our understanding of complex biological processes, offering insights into cellular heterogeneity and gene expression dynamics [1–11]. This technology enables detailed exploration of developmental trajectories, cell type-specific responses, and regulatory networks at the individual cell level. Despite its transformative impact, current single-cell RNA sequencing (scRNA-seq) protocols are inherently destructive, limiting the ability to track mRNA dynamics over time and offering only static snapshots of cellular populations.

RNA velocity methods have emerged as a powerful computational approach to reconstruct the sequence of transcriptional changes in cellular states from snapshot provided by scRNA-seq data [5, 12–19]. These techniques infer time derivative of mRNA abundance by analyzing the ratio between unspliced and spliced mRNA[12, 14]. However, they are inherently constrained by the biased and often inaccurate separation of spliced and unspliced counts in current scRNA-seq protocols [13, 20]. Additionally, the timescale of splicing is often much shorter than that of mRNA degradation, and thus cannot reliably capture longer-term transitions between cellular states.

To address these limitations, we introduce noSpliceVelo, a novel technique to infer RNA velocity that does not rely on separating unspliced and spliced transcripts. Instead, our method leverages the temporal relationship between gene expression variance and mean coming from a biophysically motivated and experimentally confirmed model of bursty gene expression [21–24]. noSpliceVelo is particularly effective for genes exhibiting significant burstiness and whose expression changes over longer timescales. Thus, it complements splicing-based RNA velocity inference techniques, which are more accurate for genes with expression changes over shorter timescales comparable to the inverse splicing rate.

We evaluate noSpliceVelo’s performance by applying it to scRNA-seq data for pancreatic endocrinogenesis [25], mouse and human erythroid maturation [26, 27], and neuronal stimulation in the embryonic mouse cortex [4], and comparing the results with those generated by scVelo [14], a popular modern splicing-based RNA velocity method. In pancreatic endocrinogenesis, noSpliceVelo performs comparably to scVelo, effectively capturing the progression of gene expression from progenitor to terminal states. For erythroid maturation in mouse and human datasets, noSpliceVelo accurately reflects the biologically correct trajectory from immature to mature cellular states, while scVelo incorrectly infers segmented and backward flows. Finally, for neuronal stimulation in the embryonic mouse cortex, the pseudotime inferred by noSpliceVelo shows a strong positive correlation (Pearson’s correlation coefficient of 0.8) with the experimental time.

Beyond its performance in trajectory inference, noSpliceVelo leverages its underlying biophysical model to infer key kinetic parameters of gene regulation: burst frequency and burst size. Burst frequency quantifies the rate at which a promoter actively transcribes mRNA, serving as an aggregate parameter for multiple upstream processes, including chromatin remodeling, transcription activator binding, and transcription initiation complex assembly [28]. For bursty or noisy genes, burst size represents the average number of mRNA molecules transcribed in short succession during a promoter’s transcriptionally active state [23, 24, 28].

To validate noSpliceVelo’s parameter estimation, we compared our computationally inferred values of burst size and burst frequency in mouse erythroid maturation to their experimental measurements in mouse embryonic stem cells, as specific measurements for the cellular states in our analyzed datasets were unavailable. The parameters estimated by noSpliceVelo showed significant correlation with these experimental measurements in a different cellular state, achieving a Pearson’s correlation coefficient of up to 0.5. We anticipate even stronger correlations when comparing to experimental measurements directly matched to cellular states in our analyzed datasets. This ability to infer biologically relevant parameters positions noSpliceVelo as a valuable tool for uncovering the underlying mechanisms driving gene expression changes.

## 2 Model and results

### Model of bursty gene expression

We begin with an experimentally verified biophysical model of bursty gene expression in single cells [22, 23, 29–32] (Fig. 1a). In this model, mRNA is transcribed from a promoter in bursts that are geometrically distributed [23] with mean size *β*. Biologically, for eukaryotic genes, the burst frequency (denoted as *f*) represents the rate at which the chromatin is in the open state and the promoter actively transcribes mRNA. Burst frequency typically results from the integration of various upstream processes, including binding of transcription activators and enhancers, and the assembly of the transcription initiation complex [28]. We assume that the waiting time between successive transcription bursts is exponentially distributed. Each mRNA molecule is assumed to be degraded at a constant rate *γ*.

**Fig. 1:**
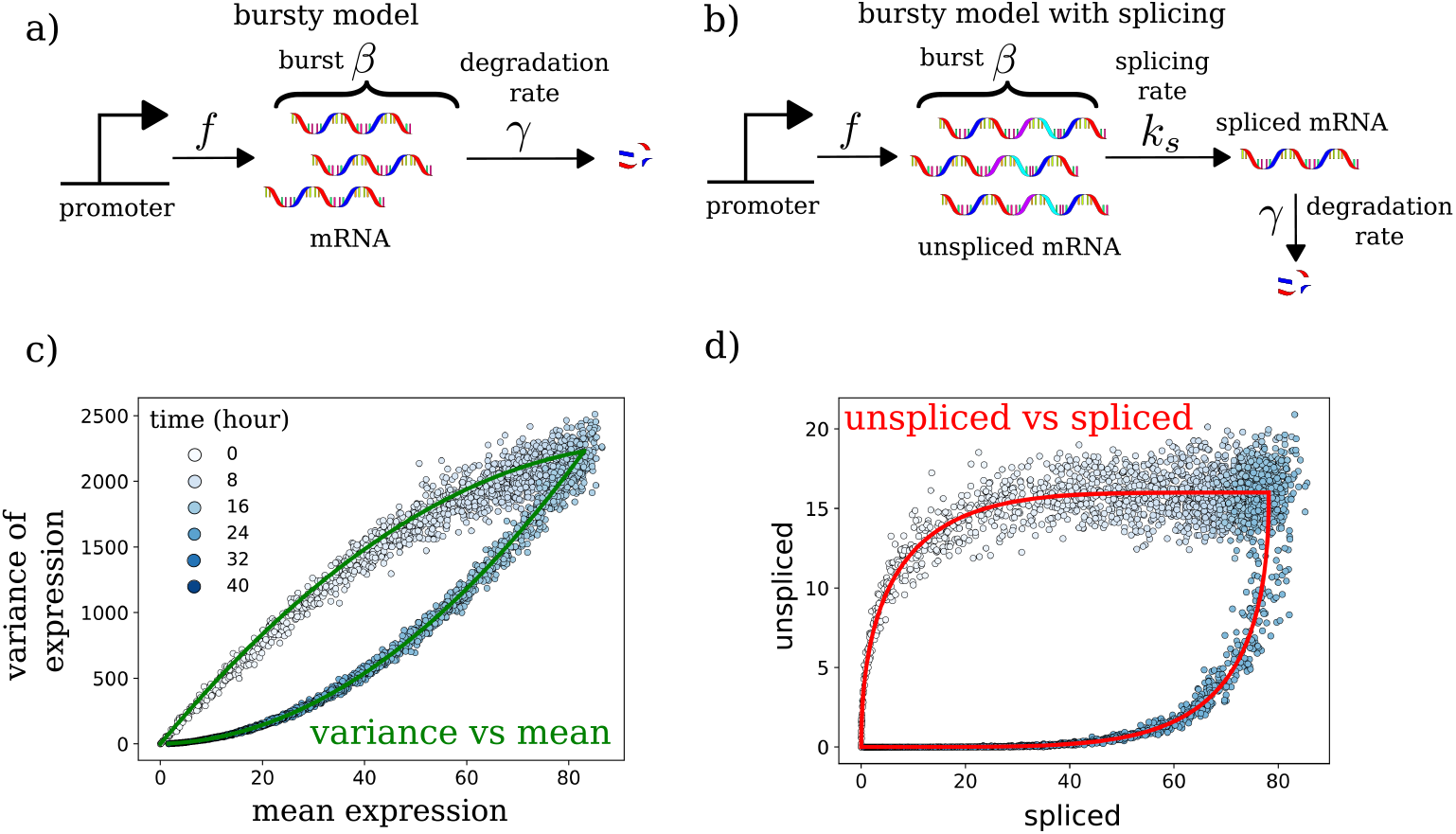
The biophysical model underlying noSpliceVelo. a) The biophysical model of bursty gene expression used in noSpliceVelo: The promoter transcribes mRNA at a rate equal to the burst frequency *f*. Each transcription event produces a geometrically-distributed burst of mRNA with an average size *β*. Each mRNA molecule is then degraded at a constant rate *γ*. b) The expanded bursty model incorporating splicing dynamics used to demonstrate the similarity between unsplicedvs-spliced and variance-vs-mean phase plots for RNA velocity inference. Unspliced mRNA is produced through the bursty process, while splicing occurs as a first-order reaction with rate *k*_*s*_, converting one unspliced mRNA into one spliced mRNA. c) Scatterplot showing the variance versus the mean of gene expression from a simulation of the expanded bursty model (panel b). The solid green curves represent parabolic fits based on the analytical solutions for this model. d) Similarly, scatterplot showing the relationship between unspliced and spliced abundances from a simulation of the expanded bursty model (panel b). The solid red curves are fits based on the analytical solutions for the model. In panels c and d, the expanded bursty model was simulated with a sudden increase in *f* from 0 at the start of the simulation. After some time, before reaching the steady state, *f* suddenly switches back to 0. In these illustrations but not in our model fits *β* is assumed to remain constant throughout the cycle of upregulation in *f* followed by its downregulation.

The temporal dynamics of the probability distribution of mRNA counts in the bursty gene expression model are governed by a chemical master equation (see Methods section, eq. (4)), which can be solved to obtain differential equations for the time dependence of the mean (*µ* (*t*)) and the variance (*σ*^2^ (*t*)) of the mRNA count:

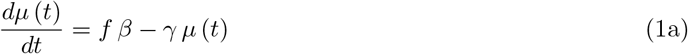

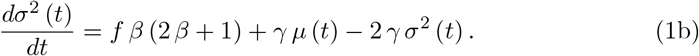

Eq. (1a) can be intuitively understood by recognizing that *f* and *β* represent promoter kinetics. Their product yields the total rate of mRNA production, which appears as the first term on the right-hand side (RHS) of the equation. The negative term on the RHS represents the total degradation rate.

In eq. (1b), we observe that *σ*^2^ (*t*) grows faster than *µ* (*t*) by a factor of 2 *β* + 1, as evident in the first term on the RHS. This difference in the rates of change between *µ*(*t*) and *σ*^2^ (*t*), which is especially pronounced for busty genes *β* ≫ 1, forms the central tenet of our method.

To obtain the steady-state values of the variance 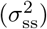 and the mean (*µ*_ss_), we set the RHS of eqs. (1)ab to zero:

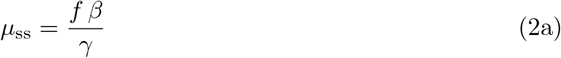

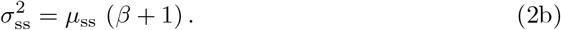

Now, consider a simple scenario where a single gene that was at a steady state abruptly changes its burst frequency *f* and/or burst size *β* at *t* = 0. Eq. (1) can be solved for *µ* (*t*) and *σ*^2^ (*t*) as follows:

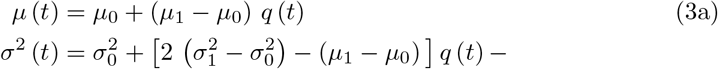

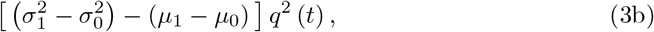

where *µ*_0_ and 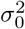 are the mean and variance of the steady state before the switch, and *µ*_1_ and 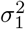 are the mean and variance of the new steady state. Here, 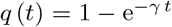 is a time-like parameter that goes from 0 to 1 as time increases from 0 to infinity.

After substituting the expression for *q* (*t*) from eq. (3a) into eq. (3b), we obtain *σ*^2^ (*t*) as a quadratic function of *µ* (*t*). The sign of the curvature in this relationship is determined by the coefficient of *q*^2^ (*t*) in eq. (3b). This coefficient tends to be negative for genes transitioning from lower to higher expression states and positive for genes decreasing their expression.

For example, when a gene’s promoter transitions from a restrictive chromatin state with *f*_0_ = *µ*_0_ = 0 to a permissive state with *f*_1_, *µ*_1_ *>* 0, the coefficient of *q*^2^ (*t*) in eq. (3b) is negative and equal to 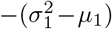. Conversely, if the gene’s promoter initially resides in a permissive chromatin state with *f*_0_, *µ*_0_ *>* 0, then switches abruptly to a restrictive state with *f*_1_ = *µ*_1_ = 0, the coefficient becomes positive and equals 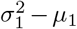.

Similar to the *σ*^2^(*t*)-vs-*µ*(*t*) phase space, the up and down transitions also have negative and positive curvatures, respectively, in the unspliced-vs-spliced phase space utilized by splicing-based RNA velocity methods. This occurs because, causally, every mRNA molecule is initially created as unspliced and subsequently processed into the spliced version. *Consequently, unspliced mRNA always leads the dynamics in the unspliced-vs-spliced phase space and the spliced mRNA follows*.

*In the case of noSpliceVelo, for bursty genes, σ*^2^(*t*) *and µ*(*t*) *follow a similar relationship, where σ*^2^(*t*) *leads and µ*(*t*) *follows*. This happens because the rate of change for *σ*^2^(*t*) is greater than *µ*(*t*) by a factor of 2 *β* + 1.

Next, consider a gene that first switches from a low to a high expression state, reaches the new high expression steady state, and then switches back to another low expression state. If the curvatures of the *σ*^2^ (*t*)-vs-*µ* (*t*) trajectories during up- and down-transitions have opposite signs (see Fig. 1b), noSpliceVelo will be able to distinguish between the two branches, thereby accurately inferring RNA velocity and time from single-cell expression data. Notably, splicing-based RNA velocity methods [12, 13] also rely on separating up from down transitions, but they use unspliced and spliced mRNA expression instead (see Fig. 1c).

Another key distinction between noSpliceVelo and splicing-based RNA velocity methods lies in different timescales of dynamical processes they rely upon. While splicing-based approaches capture dynamics on the timescale of mRNA maturation, noSpliceVelo models processes on the longer timescale of mRNA degradation. In eukaryotic cells, mRNA maturation (splicing) typically occurs within hours, whereas mRNA degradation extends over tens of hours. While shorter splicing timescales improve temporal resolution, they also lead to saturation of the spliced-to spliced ratio at longer timescales, limiting the accurate inference of RNA velocity or time beyond the inverse splicing rate. Thus noSpliceVelo should be able to achieve more accurate inference of mRNA velocity, particularly for long biological processes unfolding over tens of hours to several days.

To illustrate these analytical insights, we simulated a well-controlled biophysical model of bursty gene expression in single cells (Fig. 1b). For demonstration purposes, we extended our model to include splicing dynamics. All parameters, except for the splicing rate (*k*) and burst size *β*, were selected based on the Oct4 gene in mouse embryonic stem cells [33]. While the burst size reported in Ref. [33] was approximately 110, we reduced it to 5 to minimize biological noise in the simulation. Splicing was assumed to be five times faster than mRNA degradation. The mRNA lifetime (inverse of degradation rate) and the splicing lifetime (inverse of splicing rate) were set to 7.25 hours and 1.45 hours, respectively. Detailed simulation procedures for this example are provided in the Methods section.

We found that the up and down transitions in the simulated data within the *σ*^2^(*t*)- vs-*µ* space were well-defined by analytical parabolas with opposite curvatures (Fig. 1c). Similarly, the up and down transitions were also well-defined by analytical curves with opposite curvatures in the unspliced-vs-spliced space (Fig. 1d).

### Deep variational inference of RNA velocity without separating unspliced and spliced transcripts

The first step in our method, noSpliceVelo, is to obtain gene- and cell-specific estimates of the mean and variance of total mRNA counts. While one could theoretically estimate these values by considering a local neighborhood around each cell in a highdimensional space [12, 14, 34], such methods often introduce non-linear distortions [20]. To circumvent these issues, we directly model mRNA counts to derive accurate estimates of gene expression mean and variance.

To achieve this, we modified a popular deep variational method known as scVI [35], a variational autoencoder (VAE) [36] designed for modeling mRNA count data using the negative binomial (NB) distribution. The architecture of our modified VAE is illustrated in Fig. 2a (see Methods section for detailed modifications). We leverage the inferred gene- and cell-specific mean and variance parameters of the NB distribution as our estimates for the mean and variance of total mRNA counts.

**Fig. 2:**
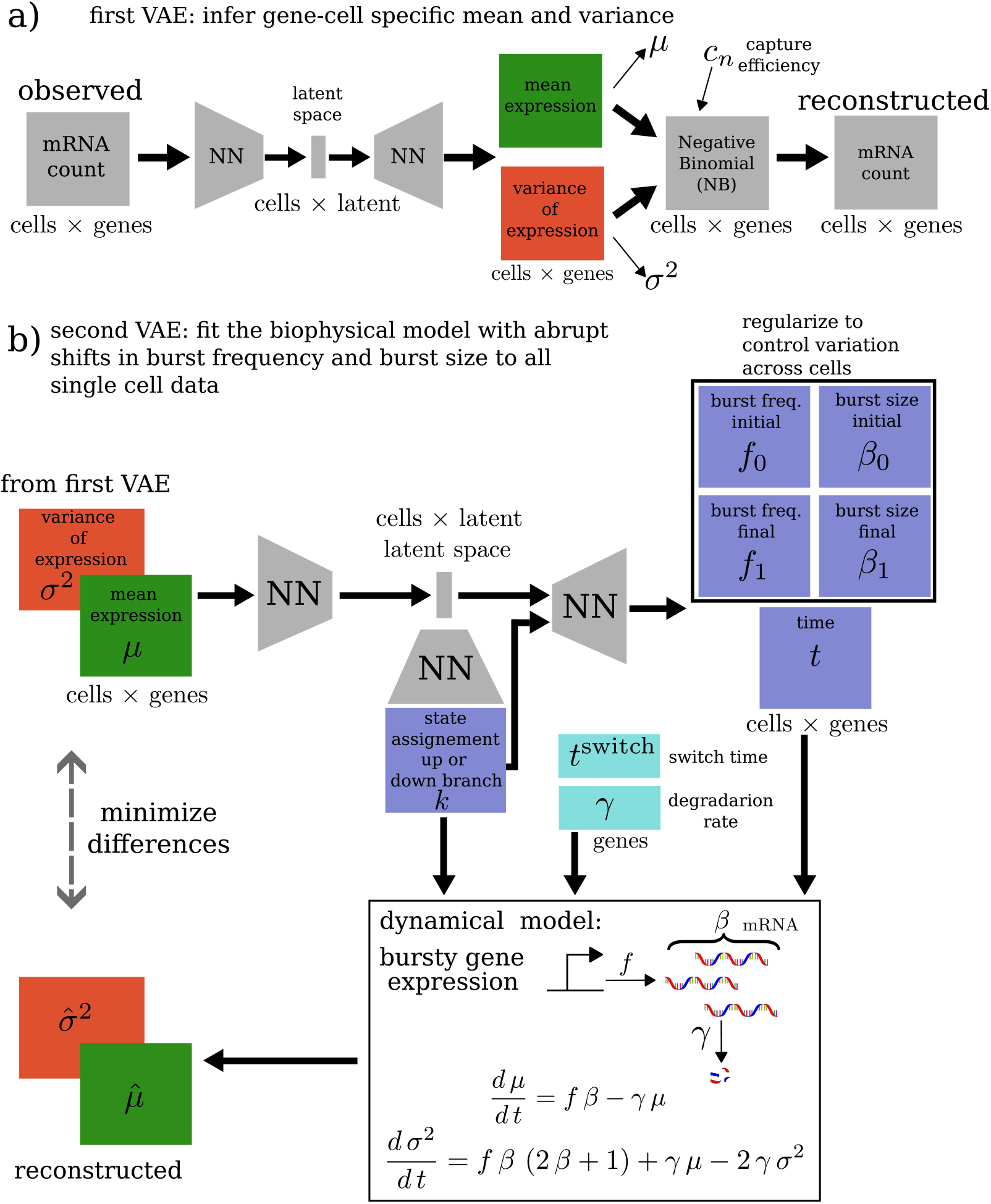
Overview of noSpliceVelo’s architecture. a) **First VAE Architecture:** This VAE estimates the mean and variance of expression from raw mRNA counts. The VAE receives gene- and cell-specific mRNA counts as input and encodes them into a latent cell representation through a neural network. The decoder network then processes this cell representation to output the mean and variance parameters for a negative binomial (NB) distribution. These parameters are used to compute the likelihood of the observed mRNA counts, and serve as estimates of the gene- and cell-specific mean and variance of expression. b) **Second VAE Architecture:** This VAE takes as input the mean and variance of expression estimated from the first VAE (shown in panel a). It encodes these estimates into a latent cellular representation, which further encodes the transcriptional state assignment (*k*) for each cell in all genes, distinguishing between upregulated and downregulated branches. The decoder network uses the latent cellular representation to generate time *t* and burst parameters for the initial (*f*_0_, *β*_0_), and the final (*f*_1_ *β*_1_) steady states. Separate neural networks decode these parameters for the upregulated and downregulated branches. However, *f*_0_, and *β*_0_ for the upregulated branch are treated as gene-specific variational parameters rather than being decoded by the network. Eq. (2) is used to convert *f*_0_ and *β*_0_ into *µ*_0_ and 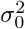, and *f*_1_ and *β*_1_ into *µ*_1_ and 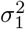. The time when genes switch from upregulation to downregulation (*t*^*switch*^) and the degradation rate (*γ*) are gene-specific variational parameters. The likelihood functions for the upregulated and downregulated branches are functions of *t, µ*_0_, 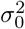, *µ*_1_, 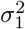, and *γ* via eq. 3.

The second stage of noSpliceVelo employs another VAE (Fig. 2b), inspired by the architecture introduced in Ref. [16]. For each cell *n*, the encoder of noSpliceVelo takes as input the gene-specific mean and variance of expression inferred from the first VAE and generates a latent cell representation *z*_*n*_ (a low-dimensional vector, with dimensionality *d* = 10 in our setup). This latent representation decodes a transcriptional state assignment variable *k*_*n*_ (with dimensionality equal to the number of genes *G*), which assigns the cells to either the upregulated or downregulated branch of gene expression. The state assignment variable *k*_*n*_ is necessary since noSpliceVelo assumes that each gene undergoes a cycle of upregulation followed by downregulation, as introduced in previous studies [12, 14, 16].

Next, *z*_*n*_ decodes the burst parameters (*f, β*) for the final steady states for both the upregulated and downregulated branches. For the initial steady state, burst parameters are only decoded for the downregulated branch, while for the upregulated branch, these are treated as gene-specific variational parameters. Although some burst parameters are allowed to be gene- and cell-specific, we impose regularization to ensure their distributions are narrow across cells. The inferred burst parameters for the upregulated and downregulated branches are converted into branch-specific *µ*_0_, 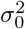, *µ*_1_, and 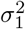 via eq. (2). Additionally, *z*_*n*_ decodes the time *t* for the two branches of gene regulation, while the mRNA degradation rate *γ* is assumed to be a gene-specific but cell-independent variational parameter.

With all the inferred parameters, *µ*(*t*) and *σ*^2^(*t*), and their corresponding likelihoods, are estimated for the upregulated and downregulated branches using eq. (3). The model is trained to minimize the evidence lower bound (ELBO) [16, 36], augmented with the above regularization for *f* and *β*. The inferred parameters can then be used to estimate the total RNA velocity, given by the right-hand side of eq. (1).

Further details on the model architecture, training process, and velocity and pseudotime estimation are provided in the Methods section.

### Application of noSpliceVelo to single cell transcriptomics data

#### Comparison between noSpliceVelo and scVelo in endocrine development in the mouse pancreas

We first demonstrate noSpliceVelo’s capabilities using real-world scRNA-seq data of endocrine development in the mouse pancreas. This dataset was chosen because it has been widely used to benchmark splicing-based RNA velocity methods [13, 15–18, 37, 38].

Following previous studies, we focus on transcriptomic profiles sampled from embryonic day (E) 15.5 [13, 25]. These profiles encompass four major terminal cell states in endocrine development: glucagon-producing *α* cells, insulin-producing *β*-cells, somatostatin-producingδ-cells, and ghrelin-producing *ϵ*-cells. Additionally, the dataset includes progenitor ductal cell states and several intermediate cell states.

In Fig. 3, we compare the performance of noSpliceVelo with scVelo [13], a popular RNA velocity method. As illustrated in Figs. 3a and 3b, the velocities estimated by noSpliceVelo for mean and variance of expression effectively capture the overall flow of gene expression. This flow depicts cells transitioning from the progenitor cell state (ductal, blue) to multiple terminal states, such as *α* (brown) and *β* (pink), via intermediate states.

**Fig. 3:**
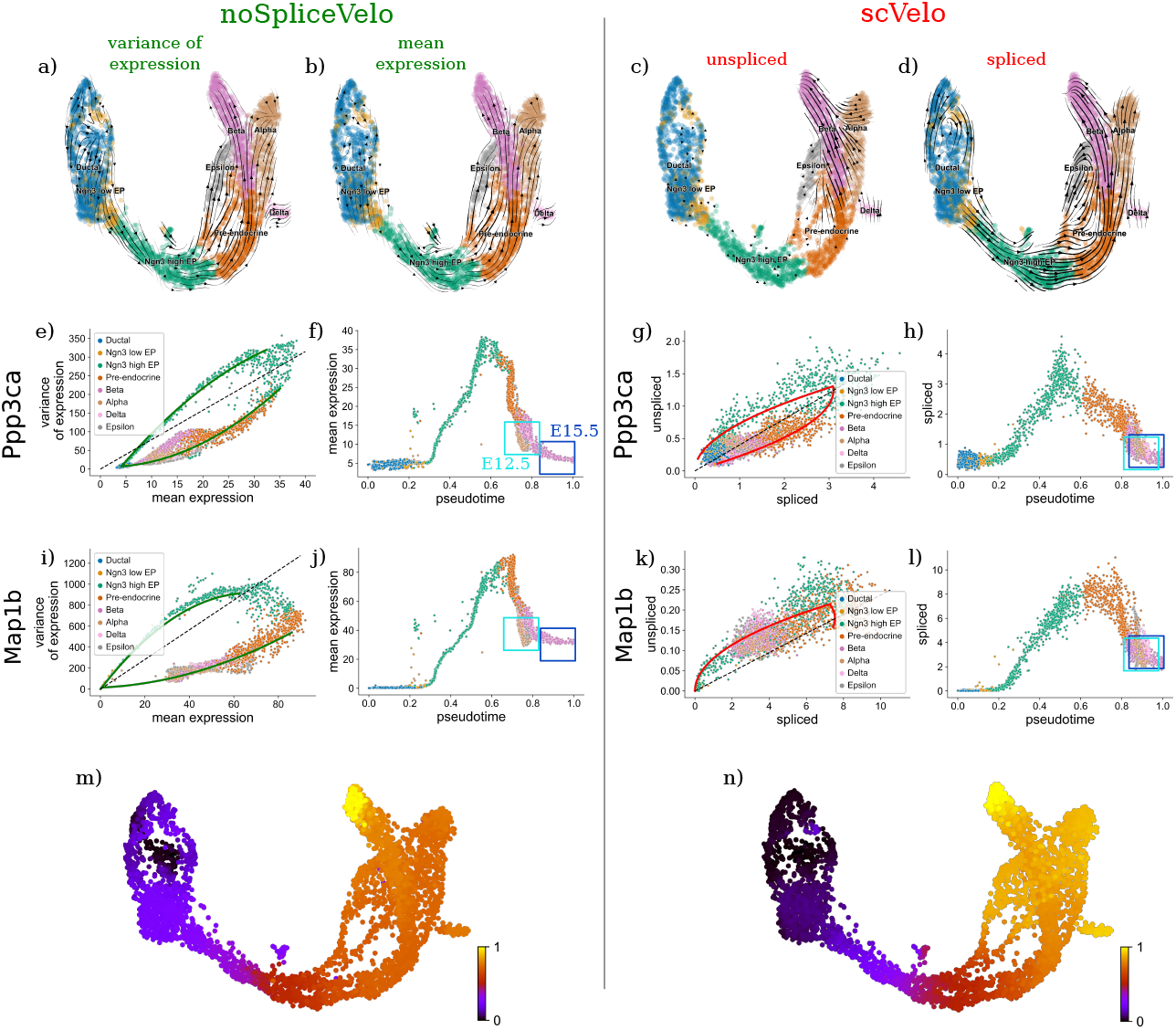
Application of noSpliceVelo to pancreatic endocrinogenesis. a-d) Projection of velocity estimates onto the UMAP representation from Ref. [14] for: a) variance of expression (noSpliceVelo); b) mean expression (noSpliceVelo); c) unspliced mRNA abundance (scVelo); d) spliced mRNA abundance (scVelo). e,i) Variance vs. mean expression phase plots for genes e) Ppp3ca and i) Map1b. Solid green curves represent parabolic fits estimated by noSpliceVelo. Dashed black lines indicate the steady state for the upregulated branch of gene expression. f,j) Mean expression for f) Ppp3ca and j) Map1b plotted against noSpliceVelo’s velocity pseudotime. Cyan and blue boxes represent *α* and *β* cells, respectively, with the embryonic days of their first appearance mentioned next to the boxes. g,k) Unspliced vs. spliced abundance phase plots for g) Ppp3ca and k) Map1b. Solid red curves represent fits estimated by scVelo. Dashed black lines indicate the steady state. h,l) Spliced abundance for h) Ppp3ca and l) Map1b plotted against scVelo’s velocity pseudotime. m,n) Heatmaps of velocity pseudotimes inferred by m) noSpliceVelo and n) scVelo, projected onto the UMAP representation from Ref. [14].

In contrast, for scVelo, only the spliced velocity accurately captures the overall flow (Fig. 3d). The unspliced velocity calculated with scVelo erroneously shows the *α* terminal state transitioning to the *β* terminal state (Fig. 3c), which is biologically implausible. Since unspliced counts are typically sparse and noisy, estimates of unspliced velocity are unreliable.

Figs. 3e-3f and 3i-3j provide a closer look at the performance of noSpliceVelo by examining two genes, Ppp3ca and Map1b, which have been highlighted in other studies [13, 14]. For Ppp3ca, the variance-vs-mean relationship is successfully explained by the parabolic curves derived in eqs. (3)a,b. Fig. 3f illustrates the mean expression of this gene in cells as a function of pseudotime, as inferred by noSpliceVelo. Figs. 3g and 3h present equivalent results for Ppp3ca as inferred by scVelo.

Notably, the unspliced-vs-spliced phase plot for Map1b in Fig. 3k exhibits a poorer separation of upper and lower branches of gene expression compared to our method (Fig. 3i). To quantify noSpliceVelo’s success and scVelo’s failure for Map1b, we examined cells with pseudotime greater than 0.6 in Fig. 3l. Based on the global pseudotime for scVelo, these cells should be in the downregulated branch. However, scVelo only assigns 2.3% of these cells to the downregulated branch, while noSpliceVelo infers 98.4% of these cells to be in the downregulated branch.

Furthermore, the pseudotime inferred by our method consistently predicts the separation of the two main terminal cellular states: *α* (box E12.5, embryonic day 12.5) and *β* (box E15.5, embryonic day 15.5) (Figs. 3f and 3j). This separation is statistically significant, with a mean difference of 0.077 in pseudotime (p-value = 1.5 × 10^−75^, one-sided Welch’s t-test). This finding aligns with the experimental observation that *α*-cells are produced earlier than *β*-cells [13, 25]. In contrast, scVelo does not exhibit a statistically significant separation of *α* and *β* cellular states in Figs. 3h and 3l (mean difference of 0.0003 in pseudotime, p-value = 0.88, one-sided Welch’s t-test).

The temporal separation between *β* and *α* cells is also evident in the pseudotime heatmap for noSpliceVelo (Fig. 3m). Notably, our pseudotime provides a sharper contrast between the *β* and *α* states compared to scVelo (Fig. 3n). This distinction is further illustrated in the scatterplot comparing the pseudotimes for noSpliceVelo and scVelo (Supplementary Fig. (S1)).

#### Application of noSpliceVelo to mouse and human erythroid maturation

We next applied noSpliceVelo to scRNA-seq datasets of mouse [26] and human [27] erythroid maturation (Fig. 4). In Fig. 4a, noSpliceVelo accurately infers the overall trajectory of gene expression in mouse cells, starting from the immature Blood progenitor 1 state and progressing to the mature Erythroid 3 state [26, 39]. The trajectory reveals transitions through three intermediate states: Blood progenitor 2, Erythroid 1, and Erythroid 2.

**Fig. 4:**
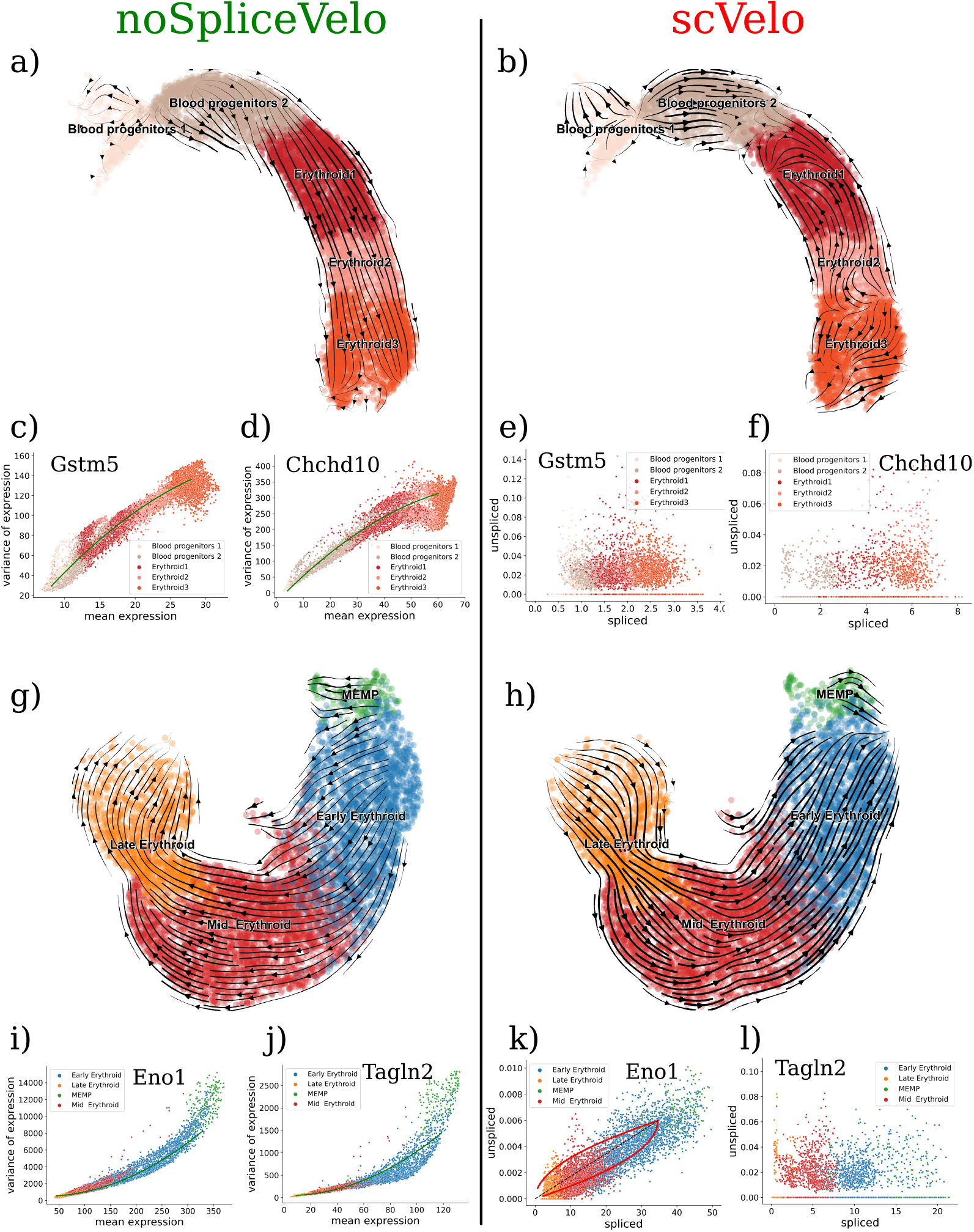
Application of noSpliceVelo to mouse and human erythroid maturation. Mouse gastrulation erythropoiesis [26]–. a,b) Projection of velocity estimates onto the UMAP representation from Ref. [13] for: a) mean expression (noSpliceVelo), and b) spliced mRNA abundance (scVelo). c,d) Variance vs. mean expression phase plots for genes c) Gstm5 and d) Chchd10. Solid green curves represent parabolic fits estimated by noSpliceVelo. e,f) Unspliced vs. spliced abundance phase plots for e) Gstm5 and f) Chchd10. Solid red curves represent fits estimated by scVelo. **Human erythroid haematopoiesis** [27]– g,h) Projection of velocity estimates onto the UMAP representation from Ref. [18] for: g) mean expression (noSpliceVelo), and h) spliced mRNA abundance (scVelo). i,j) Variance vs. mean expression phase plots for genes i) Eno1 and j) Tagln2. Solid green curves represent parabolic fits estimated by noSpliceVelo. k,l) Unspliced vs. spliced abundance phase plots for k) Eno1 and l) Tagln2. Solid red curves represent fits estimated by scVelo.

Similarly, in Fig. 4g, noSpliceVelo accurately identifies the developmental progression of human cells, beginning with the MEMP (Megakaryocyte-Erythroid-Mast cell Progenitor) state and culminating in the Late Erythroid state [27, 39]. The analysis reveals a progression through two key intermediate stages: Early and Mid Erythroid.

In contrast, Figs. 4b and 4h show that scVelo infers either backward flows or distorted trajectories for both datasets. For the mouse dataset, scVelo predicts a segmented velocity stream (Fig. 4b), inaccurately suggesting a transition from the mature Erythroid 3 state to the intermediate Erythroid 1 state via the Erythroid 2 state. In the human dataset, scVelo erroneously infers a reversed trajectory, suggesting movement from the terminal Late Erythroid state back to the immature MEMP state (Fig. 4h).

noSpliceVelo’s accurate inference of the erythroid maturation trajectory can be contrasted with scVelo’s incorrect inference of reversed transitions from mature to immature states by comparing the inferred pseudotimes, as shown in Fig. (S2)a. The pseudotimes exhibit a strong negative correlation (Fig. (S2)a). Additionally, the scatterplot between scVelo’s and noSpliceVelo’s pseudotimes displays an inverse sigmoidal shape, indicating that scVelo struggles to temporally separate cells at the beginning and end of the differentiation trajectory (Fig. (S2)a).

We further evaluated the performance of noSpliceVelo by focusing on specific genes in both the mouse and human datasets. For the mouse dataset, we analyzed Gstm5 and Chchd10 (Figs. 4c, 4d, 4e, and 4f), and for the human dataset, we examined Eno1 and Tagln2 (Figs. 4i, 4j, 4k, and 4l). Figs. 4c, 4d, 4i, and 4j present the variance-versus-mean phase plots for these genes. We observed that the parabolas fitted by noSpliceVelo (solid green curves) effectively capture the general trends in gene expression. Specifically, noSpliceVelo accurately identifies Gstm5 and Chchd10 as upregulated (on the up branch), while Eno1 and Tagln2 are correctly classified as downregulated (on the down branch).

In contrast, scVelo’s performance for these genes is significantly inferior. For example, scVelo fails to accurately model the data for Gstm5 (Fig. 4e), Chchd10 (Fig. 4f) and Tagln2 (Fig. 4l) due to insufficient correlation between unspliced and spliced RNA abundances. Additionally, the fit provided by scVelo for Eno1 (Fig. 4k) is poor and does not clearly classify this gene as downregulated.

To quantify noSpliceVelo’s ability to identify upregulation and downregulation, we calculated the number of cells with correctly inferred velocity direction. For noSpliceVelo, we generated 100 velocity samples from the posterior distribution and computed the percentage of cells with the correct direction by retaining only the sign of the velocity. For scVelo, we calculated this percentage as a point estimate since it does not provide samples from a posterior distribution of the learned parameters.

noSpliceVelo successfully predicts the correct direction for over 90% of the cells for both the upregulation of Gstm5 and the downregulation of Eno1. In contrast, scVelo fails to fit Gstm5 and predicts the correct direction for only about 70% of the cells for Eno1.

#### noSpliceVelo accurately captures short-term and long-term activation in activity-induced neurons

We next applied noSpliceVelo to scRNA-seq data from neurons in the embryonic mouse cortex [4]. In this study, mouse cortical cultures were metabolically labeled and stimulated for varying durations of neuronal activity (0, 15, 30, 60, and 120 minutes). Cells were sampled at each timepoint and sequenced together. We assumed that this collection of cells approximates a stimulated population where cells were sampled at different timepoints after introduction of the stimulation. We utilized only the total mRNA count, discarded any information about metabolic labeling, and retained only neural activity genes [4, 6] for analysis with noSpliceVelo.

We projected noSpliceVelo’s inferred velocity for mean expression onto the UMAP-based low-dimensional representation of its latent space. The resulting velocity streams, shown in Fig. 5a, accurately reflect the transition of neurons from 0 to 15, 30, 60, and 120 minutes. Additionally, Fig. 5b demonstrates a strong positive correlation between noSpliceVelo’s inferred pseudotime and experimental time (Spearman’s rank correlation of 0.8, p-value = 1 × 10^−323^). The pseudotimes for any two successive experimental timepoints were also statistically significantly different based on one-sided Welch’s t-tests (Fig. 5b).

**Fig. 5:**
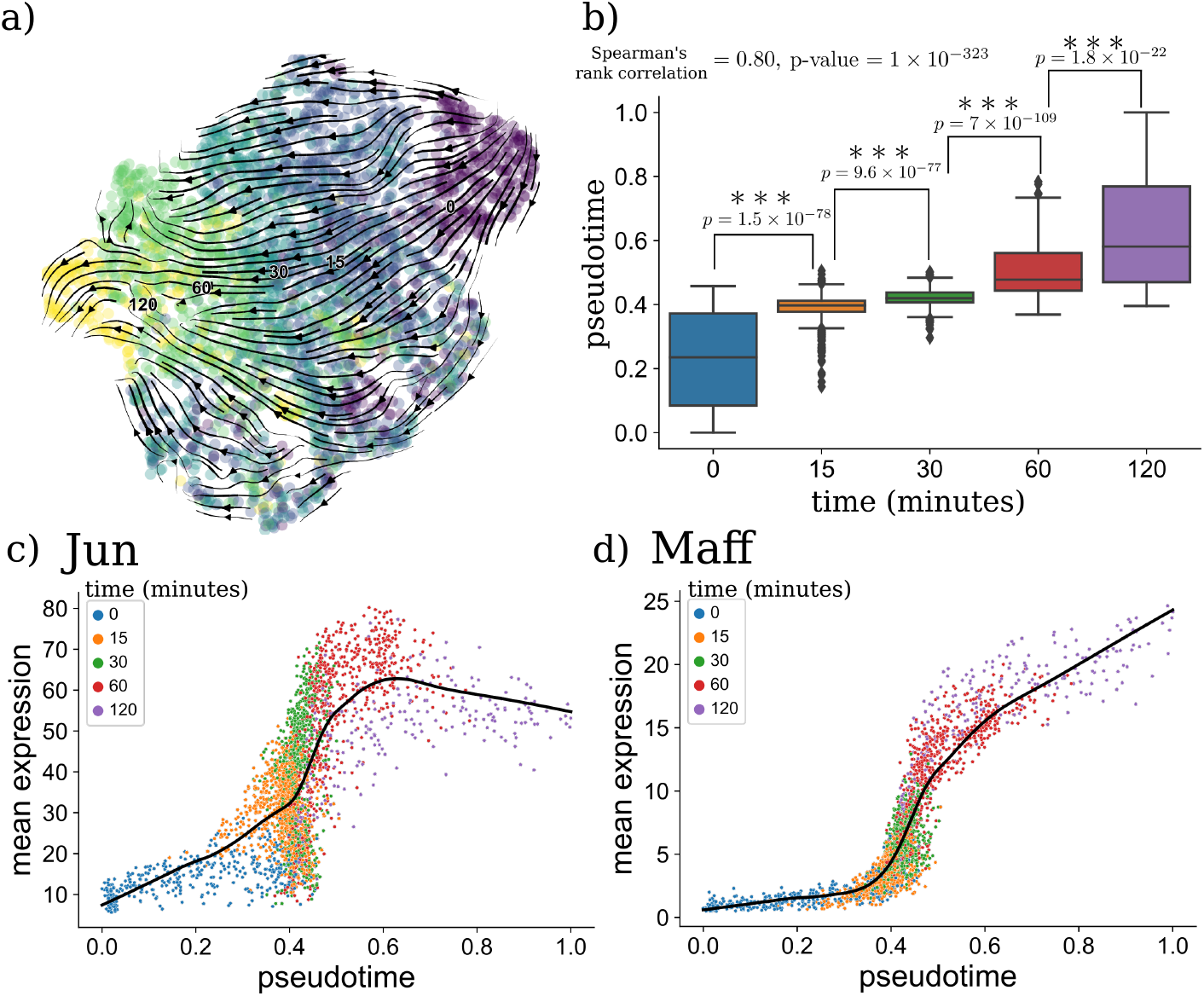
noSpliceVelo correctly infers temporal ordering and activation patterns in embryonic mouse cortex. a) Velocity estimated by noSpliceVelo for mean expression, projected onto the UMAP representation for noSpliceVelo’s latent space. The projection correctly identifies the flow of cells from 0 to 15, 30, 60, and 120 minutes. b) noSpliceVelo’s velocity pseudotime, y-axis, shows strong positive correlation with experimental time, x-axis, (Spearman’s rank correlation of 0.8, p-value = 1 × 10^−323^). Pseudotimes for successive experimental timepoints are statistically significantly different (one-sided Welch’s t-tests; p-values shown on connecting bars). c,d) Mean expression plotted as a function of noSpliceVelo’s pseudotime for the earlyresponse gene c) Jun and the late-response gene d) Maff. Solid black curves represent LOWESS (locally weighted scatterplot smoothing) regression fits.

The pseudotime inferred by noSpliceVelo successfully captured both short- and long-term activation patterns in the dataset. This is evident from the mean expression vs. pseudotime plots for the early response gene Jun (Fig. 5c) and the late response gene Maff (Fig. 5d). The mean expression for Jun peaks around a pseudotime of 0.6, which corresponds to the scatter of cells at 60 minutes of experimental time. Thereafter, the mean expression for Jun declines, consistent with previous observations [4]. In contrast, the late response gene Maff shows a monotonic increase in mean expression as a function of pseudotime, consistent with previous findings [4].

## Discussion

We have developed noSpliceVelo, a novel method for inferring total mRNA velocity from scRNA-seq data without relying on the separation of spliced and unspliced mRNA transcripts. This approach is advantageous in scenarios where splicing data is difficult to obtain (such as with short-read sequencing) or of low quality. noSpliceVelo addresses the limitations of splicing-based RNA velocity methods, which can be unreliable when scRNA-seq data is dominated by spliced mRNA counts with sparse and unreliable unspliced counts. Importantly, noSpliceVelo captures dynamics on the timescale of mRNA degradation (tens of hours to days), making it suitable for studying dynamic processes such as development and differentiation. This contrasts with splicing-based methods, which typically model dynamics on much shorter timescales (minutes to several hours).

Our results demonstrate that noSpliceVelo performs comparably to, and in some cases outperforms, splicing-based methods like scVelo. In the context of erythroid maturation in both mouse and human, noSpliceVelo accurately captures the correct direction of differentiation from immature progenitor to mature terminal states, including transitions through intermediate stages. In contrast, scVelo provides a segmented velocity flow with some segments exhibiting opposing directions, and its velocity pseudotime incorrectly suggests gene expression flows from mature to intermediate or immature states.

One of the reasons for noSpliceVelo’s robustness is its approach to estimating input mean and variance of expression via direct mRNA count modeling. This method addresses issues commonly associated with techniques that rely on ad-hoc normalization of counts. As discussed in Ref. [20], such normalization techniques can introduce artifacts, making our method’s approach more robust and reliable. Our method then interprets the mean and variance of mRNA copy numbers in individual cells using a standard biophysical model of bursty gene expression (Refs. [21–23]). In contrast, most splicing-based RNA velocity methods employ a simpler constitutive model of gene expression that does not account for burstiness (see Ref. [20]).

After an abrupt increase in burst frequency and/or burst size of a gene, the variance of its expression will initially rise more quickly than the mean expression. Conversely, if there is a sudden decrease in burst frequency or size, the variance will decrease more rapidly than the mean. Thus, in both cases, the variance of expression leads the dynamic change, while the mean expression follows. This is analogous to the relationship between unspliced and spliced mRNA levels, where the unspliced expression leads the sudden change in gene expression and the spliced expression follows. noSpliceVelo leverages the curvature of the variance-vs-mean plot to separate the instances where gene expression increases (negative curvature) from the cases where gene expression goes down (positive curvature).

Using the biophysical model, our method infers key kinetic parameters of gene regulation, specifically burst frequency and burst size. Since experimental measurements for these parameters are unavailable for the cell states in our analyzed datasets, we compared noSpliceVelo’s inferred burst frequency and burst size with the closest available experimental measurements from a genetically homogeneous population of mouse embryonic stem cells (mESCs) [40]. Our predicted burst sizes showed a positive correlation with experimental measurements (Pearson’s correlation coefficient (PCC) = 0.24, p-value = 3.3×10^−5^). Similarly, burst frequency also exhibited a positive correlation (PCC = 0.49, p-value = 3.3 × 10^−19^). We anticipate that these correlations would be even stronger if experimental measurements were directly matched to the dynamic cellular states observed in our analyzed datasets. Although biophysical parameters are expected to vary across different cellular contexts, this comparison allows us to assess the similarity in burst frequency and burst size between different tissue types.

In order to further enhance the accuracy and applicability of the noSpliceVelo method, in the future we plan to modify it along the following directions:

- Incorporate Splicing Dynamics: Another potential extension for noSpliceVelo is to incorporate splicing dynamics, which are currently not considered in the method. This enhanced version could simultaneously infer RNA velocity using both the ratio between unspliced and spliced mRNA, as done in scVelo, and the variance-vs-mean analysis employed by noSpliceVelo. Integrating these two dynamic processes, each with distinct time scales, is expected to enhance the overall accuracy of the method.
- Separate Regularization Terms: Although noSpliceVelo currently permits burst size and burst frequency to vary between individual cells, the regularization term encourages these distributions to remain narrow. Future work could involve modifying our method to incorporate separate regularization terms for different cellular states. This adjustment would allow burst size and burst frequency to vary between states while maintaining consistency within cells from the same state.
- Analyze Changes in Burst Dynamics: Most splicing-based RNA velocity methods typically analyze the ratio of gene expression levels in high and low expression states. In contrast, our method decomposes this ratio into a product of two distinct ratios: changes in burst size and burst frequency. Analyzing these changes could provide insights into the mechanisms driving gene expression alterations, potentially distinguishing between signatures of transcriptional regulation and epigenetic modifications [30, 41].
- Comprehensive Biophysical Model: Eukaryotic gene expression is affected by several distinct dynamic processes, including the opening and closing of chromatin by epigenetic regulation, transcription of mRNA from open chromatin influenced by transcription factors binding to promoters and enhancers, and mRNA splicing and degradation. These first two processes are known to affect both burst frequency and burst size [31, 42–46] used in our model. In future work, we aim to develop a comprehensive “soup-to-nuts” biophysical model of gene expression. This model will begin by using scATAC-seq data to infer chromatin dynamics, similar to the recently proposed ‘chromatin velocity’ method [47] and biophysical model for chromatin dynamics [48]. Chromatin dynamics inferred from scATAC-seq data will be combined with scRNA-seq data collected from the same cell. For this, we will integrate the noSpliceVelo algorithm with known gene regulatory networks, allowing for the joint inference of RNA velocities for transcription factors and their target genes, rather than assuming their independence as we currently do. Additionally, the model will incorporate mRNA splicing and degradation dynamics. By combining these mechanisms, we expect to significantly improve RNA velocity estimation and the inference of cellular state transitions. The enhanced model could also iteratively predict new edges in gene regulatory networks and uncover previously unknown epigenetic modifications.

## Methods

### Chemical master equation for the bursty model of gene expression

The chemical master equation for the bursty model in Fig. 1a is

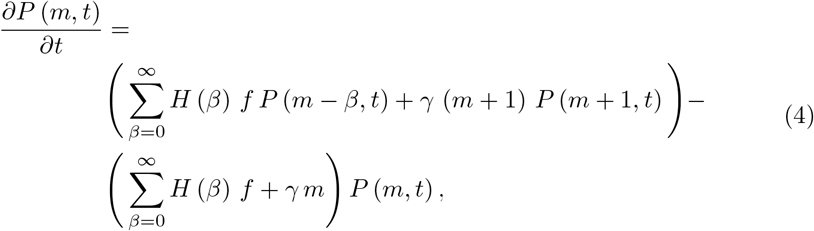

where *m* is the mRNA count, *P* (*m, t*) is the time-dependent probability distribution for *m*, and *H* (*β*) is the probability distribution for the burst size *β*, which are expected to be the geometrically distributed [29]. Eq. (4) describes a birth-death process, where the positive and the negative terms on the right hand side (RHS) come from the birth and death processes for mRNA, respectively. Temporal evolution of mean expression, *µ*(*t*), in eq. (1a) can be obtained by multiplying eq. (4) with *m* and then taking expectation on both sides. Time dependence of ⟨*m*^2^⟩ can also be obtained in a similar fashion by multiplying eq. (4) with *m*^2^ and then taking expectation on both sides. Collectively, time dependence of mRNA variance in eq. (1b) is obtained from 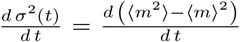. Eq. (1) was derived with the assumption that *H* is the geometric distribution. Consequently, we assume that *β* represents the average burst size in Eq. (1).

### Simulation for the expanded biophysical bursty model with splicing

We simulated the expanded bursty model with splicing in Fig. 1b for 100 identical, but independent genes across multiple cells. For each gene, the average burst size *β* was constant, while the burst frequency made up and down transitions as described previously. Each gene was simulated in every cell to go through the cycle of up and down transitions independently. 2500 cells were selected in both the up and down transitions. Finally, we had an mRNA count matrix with the 100 genes sampled across 5000 cells. The variance-vs-mean and the unspliced-vs-spliced phase plots averaged acrossed the 100 genes are shown in Figs. 1c, and 1d, respectively. The opposite curvatures in the up and down transitions are clearly visible in both the phase spaces. We also see that unspliced mRNA saturates quickly to the steady state (Fig. 1d). Whereas, variances grows/decays gradually towards steady state in Fig. 1c.

### Estimating mean and variance of expression via mRNA count modeling

We start with describing the estimation procedure for the inputs to noSpliceVelo, namely the mean and variance of expression. They are obtained by modeling raw mRNA count data with a VAE similar to scVI [35] (Fig. 2a). For approximating technical errors due to capture efficiency and sequencing depth, scVI either uses the observed library size or estimates it during training of the model. However, we use a binomial measurement model with a cell-specific capture efficiency *c*_*n*_ [49–52]. Importantly, the binomial model can explain the statistics of excess zeros seen in scRNA-seq without resorting to zero-inflation models [50, 52], which are not biophysically motivated. Furthermore, the binomial model is intuitive since all the transcripts in a given cell will be roughly captured with the same probability *c*_*n*_.

Assume that there are *N* cells and *G* genes in the observed scRNA-seq count matrix *Y*. Under the binomial measurement model the probability of observing *y*_*ng*_ transcripts of gene *g* in cell *n* is given by:

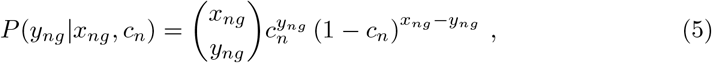

where *x*_*ng*_ is the true count of transcripts of gene *g* in cell *n*, which is unobserved, and *c*_*n*_ is the probability of sampling a single molecule of mRNA transcript, irrespective of the gene, in cell *n*.

For the expression model, we model transcript counts using the negative binomial distribution, which has biophsyical motivation [23, 24, 29]:

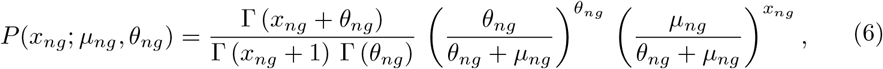

where *µ*_*ng*_ and *θ*_*ng*_ are the geneand cell-specific mean and inverse-dispersion for the negative binomial distribution. For the bursty model of gene expression (Fig. 1 a), distribution of mRNA counts at steady state is negative binomial [22, 23]. Furthermore, distributions for dynamical processes can also be well approximated with the negative binomial distribution [29].

From Eqs. (5) and (6), marginal distribution for the observed transcript count, *y*_*ng*_, is negative binomial with *c*_*n*_ *µ*_*ng*_ and *θ*_*ng*_ as the mean and the inverse-dispersion, respectively:

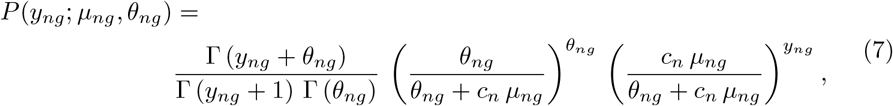

The observed mean and variance are then modeled by the mean and variance obtained from eq. (7),

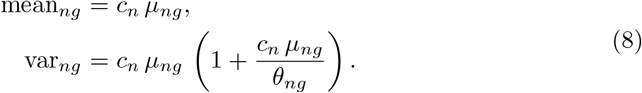

Eq. (8) represents the estimates of mean and variance of expression for the observed mRNA counts, whereas *µ*_*ng*_ and *θ*_*ng*_ are the mean and variance for the true, unobserved counts. *µ*_*ng*_ and *θ*_*ng*_ are used as inputs for noSpliceVelo’s second VAE in Fig. 2b. *µ*_*ng*_ and *θ*_*ng*_ are estimated from the first VAE (Fig. 2a) after training. *c*_*n*_ is approximated using the 2000 least variable genes (LVG) from the raw count matrix *Y* using the expression given in ref. [51]:

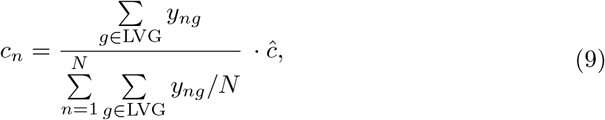

where *ĉ* is the average capture efficiency in the scRNA-seq protocol. We use *ĉ* = 0.1 in all our experiments.

The first VAE was trained similar to scVI [35] using the same optimizer and set of hyperparameters.

### noSpliceVelo model specification

#### Model assumptions

We assume that at time *t* = 0 any specified gene is first upregulated with abrupt changes in burst frequency and/or burst size. The gene may or may not reach a steady-state before it switches to the downregulated branch at some switching time 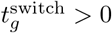. Downregulation is also accompanied by abrupt changes in burst frequency and/or burst size. Further, to ensure identifiability [53], we also assume that all genes are on the same time scale with maximum time set to *t*_max_ = 24.

#### noSpliceVelo generative process

noSpliceVelo’s generative process is similar to that introduced in VeloVI [16], a VAE for inferring splicing-based RNA velocity.

First, we sample the latent cellular representation *z*_*n*_ for each cell from a 10− dimensional unit Normal distribution. Next, for each gene *g* in cell *n*, we sample state assignments *k*_*ng*_ (0 for upregulation and 1 for downregulation) from a categorical distribution with equal probabilities for the two states.

In Eq. (3), *t* represents the time from the initial state. For upregulation, 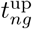 is the absolute time. For downregulation, 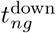 is relative to the switching time 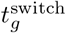, making the absolute time 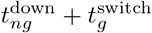. The maximum values for 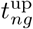 and 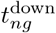 are 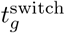 and 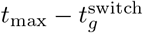, respectively.

For each cell *n*, these times are obtained by decoding *z*_*n*_ into *G*-dimensional vectors via fully connected neural networks. Softplus activation with output clamping ensures the maximum time constraints are satisfied:

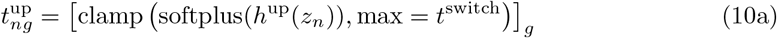

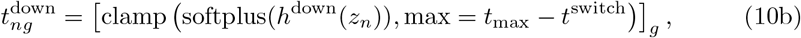

where *h*^up^ : ℝ^*d*^ → ℝ^*G*^ and *h*^down^ : ℝ^*d*^ → ℝ^*G*^ are parameterized as fully connected neural networks.

Similar to *t*, burst parameters for the initial steady state (*f*_0_ and *β*_0_) and the final steady state (*f*_1_ and *β*_1_) are decoded by fully connected neural networks via the following relationships:

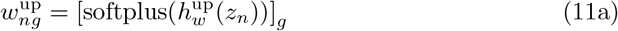

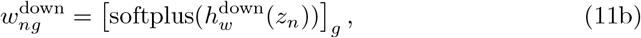

where *w* ∈ {*f*_0_, *β*_0_, *f*_1_, *β*_1_} for eq. (11b), but *w* ∈ {*f*_1_, *β*_1_} for eq. (11b).

We also impose the following constraints on the burst parameters to ensure that the upregulation and downregulation curves are monotonic in the variance-vs-mean phase plot:

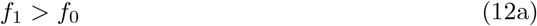

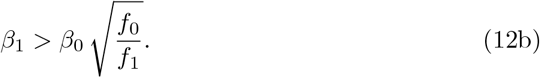

Next, eq. (2) is used to convert *f*_0_ and *β*_0_ into *µ*_0_ and 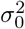, and *f*_1_ and *β*_1_ into *µ*_1_ and 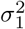. Separate values are generated for the upregulated and downregulated branches.

The mean expression is generated by sampling from a normal distribution, with the mean of this distribution given by eq. (3a):

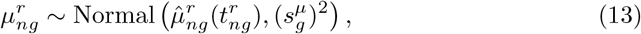

where 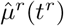 is given by eq. (3a) with the parameters inferred by the decoder, *r* ∈ {up, down}, and *s*^*µ*^ is a gene-specific scale parameter for *µ*, estimated as a variational parameter during training.

We do not directly sample the variance of expression, but rather its sqaure root (standard deviation of expression) from a normal distribution, with the mean of the distribution given by the square root of eq. (3b):

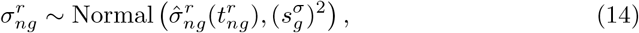

where 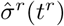 is given by the square root of eq. (3b) with the parameters inferred by the decoder, *r* ∈ {up, down}, and *s*^*σ*^ is a gene-specific scale parameter for *σ*, estimated as a variational parameter during training.

#### noSpliceVelo inference procedure

Similar to Ref. [16], we use variational inference to obtain: 1) posterior probability distributions for the latent variables, 2) point estimates of the degradation rate and the switching time, and 3) point estimates of the parameters of the neural networks. After inference, velocity can be calculated as a functional of the variational posterior.

#### Variational posterior for noSpliceVelo

We assume that the joint posterior distribution for *z* and *k* can be factorized as

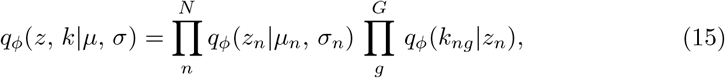

where 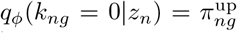 and 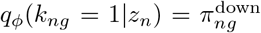, with 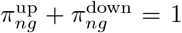. In eq. (15), dependencies are modeled using neural networks with parameter set *ϕ*. Note that *z* factorizes over all cells and *k* over all cells and genes.

Empirically, we found that working with the standard deviation of expression, rather than the variance directly, allows noSpliceVelo to accurately separate the upregulated and downregulated branches of gene expression. Therefore, the model likelihoods are computed for the mean and the standard deviation of expression. These likelihoods are given by

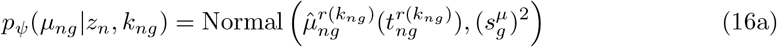

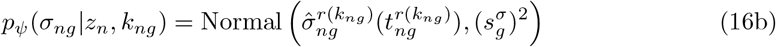

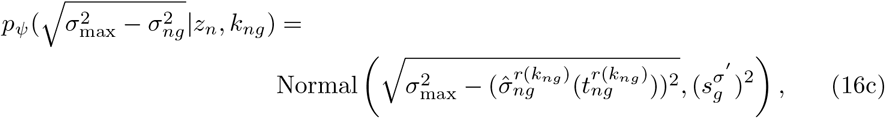

where *ψ* is the set of generative parameters (*γ*, switch time, and neural network parameters), *r*(*k*_*ng*_ = 0) = up, and *r*(*k*_*ng*_ = 1) = down. Empirically, we observed that eq. (16c) was required for noSpliceVelo to accurately separate the up and down branches of gene expression. Consequently, the variance of expression input to noSpliceVelo in Fig. 2b is first non-linearly transformed into the standard deviation of expression and 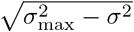, and both are used as separate inputs to the VAE. 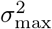 is a gene-specific, pre-determined value to ensure that the terms inside the square root are positive. We set 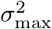 equal to five times the maximum value of variance of expression.

#### Loss function

The loss function to be minimized is defined as

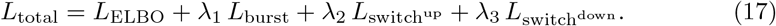

*L*_ELBO_ is the negative evidence lower bound [54] expressed as:

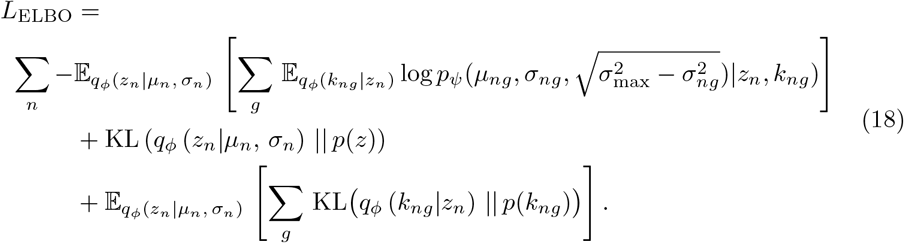

This can be estimated using randomly sampled minibatches of the data.

*L*_burst_ is a regularization term ensuring narrow distributions of burst frequency and size across cells. To do this, we include gene-specific variational parameters for 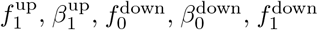, and 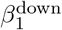 in the model. To ensure that *L*_burst_ is on the same scale as *L*_ELBO_, we compare mean and variance rather than burst parameters directly, converting them using equations (2)a,b. Then, the burst parameters are regularized by minimizing the difference between gene- and cell-specific means (variances) decoded by the latent cellular space and gene-specific means (variances). *L*_burst_ comprises six regularization terms:

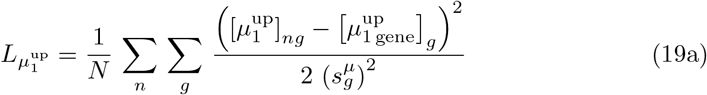

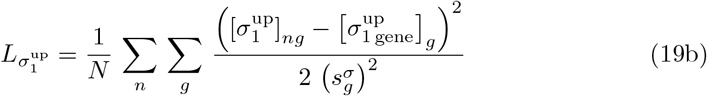

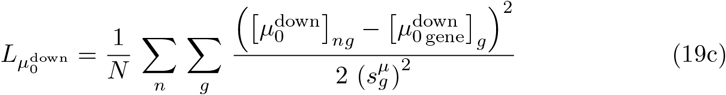

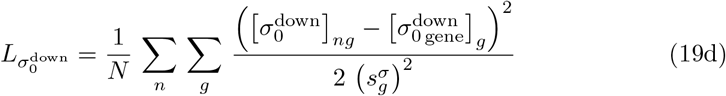

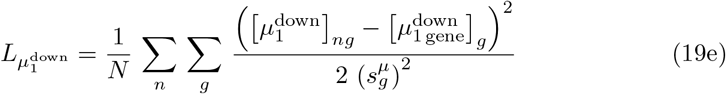

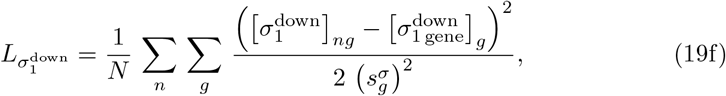

where terms with ‘gene’ in the subscript are derived from gene-specific variational burst parameters via eqs. (2)a,b. Thus, 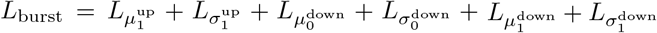.

For the penalty term 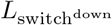, we identify cells above the 95th percentile of mean expression and compute their average mean and variance of expression, denoted as 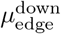, and 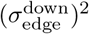. Then, the penalty term is:

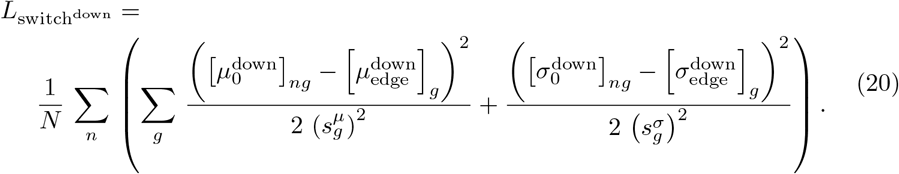

Similarly, for the penalty term 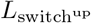, we identify cells that are above the 95the percentile of variance of expression, calculate their average mean 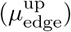 and variance 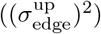 of expression. We also estimate the maximum possible values of mean and variance of expression for the upregulated branch at the switch time 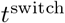, and denote them as 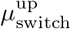 and 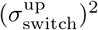. Now, 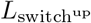 is given by

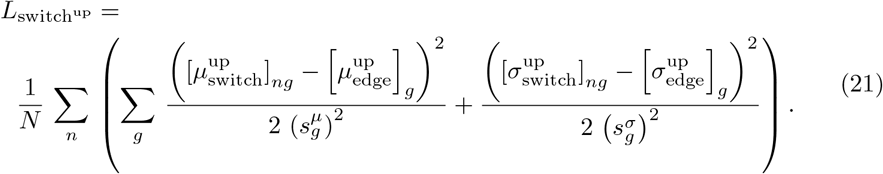

In all experiments, we set *λ*_1_ = *λ*_2_ = *λ*_3_ = 0.1.

#### Training noSpliceVelo

We optimized noSpliceVelo using the same settings and default parameters as described in Ref. [16]. All neural networks are fully connected, with ReLU activation functions for hidden layers and softplus activation for non-negative distributional parameters.

#### Downstream tasks

For all downstream inferences, such as estimating fitted values for mean and variance of expression and predicting RNA velocity, we employ posterior predictive inference. The general steps are as follows:

1. Sample *z*_*n*_ from *q*_*ϕ*_ (*z*_*n*_|*µ*_*n*_, *σ*_*n*_), and estimate the probability distribution *q*_*ϕ*_ (*k*_*ng*_|*z*_*n*_) based on the sample *z*_*n*_
2. Identify the state 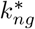 with the highest probability 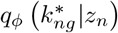.
3. Compute 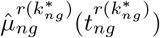 and 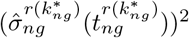 from eqs. (3a) and (3b), where 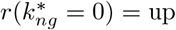 and 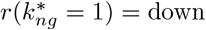. Also, calculate the velocity for mean and variance of expression using equations (1a) and (1b) with the inferred parameters for the selected state 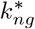.
4. Repeat steps 1 to 3, 100 times, and average the values of 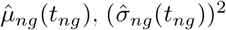, and the velocities across these repetitions.

We select the state with the highest probability in step 2 and do not average across states because velocities in the upregulated and downregulated branches can differ in scale, and averaging may lead to inaccurate velocity direction assignments.

Velocity pseudotime was estimated from the inferred velocities using the method introduced in [14].

#### Data prepreprocessing

The following steps were used to preprocess all datasets:

1. Exclude cells with non-zero mRNA counts in fewer than 100 genes from further analysis.
2. Estimate capture efficiency from the 2000 least variable genes based on dispersion, as described in equation (9).
3. Select the top 2000 highly variable genes based on dispersion [55] for further analysis.
4. Model raw count data using our first VAE (Fig. 2b) to obtain mean and variance of gene expression.
5. Smooth the obtained estimates of mean and variance of gene expression using a nearest neighbor graph with *k* = 30 neighbors in the latent space of the first VAE.

Some preprocessing steps were dataset-dependent:

1. For the human erythroid maturation dataset [27], there were roughly 36, 000 single cells. We randomly subsampled this dataset to retain 25% of the cells for our analysis.
2. For the mouse embryonic cortex dataset [4], following previous studies [4, 6], we retained only the neural activity genes for analysis.

## Data availability

All the datasets analyzed in the paper are publicly available and published. The processed and annotated data for mouse pancreatic endocrinogenesis and mouse gastrulation erythropoiesis are available with the Python package for scVelo at https://scvelo.org. For human erythroid differentiation, the processed and annotated dataset is available with a splicing-based RNA velocity method UniTVelo at https://github.com/StatBiomed/UniTVelo/tree/main. Finally, the dataset for emrbyonic mouse cortex is available with the Python package Dynamo at https://dynamo-release.readthedocs.io/en/latest/index.html.

## Code availability

All the results reported in this paper and our Python implementation of noSpliceVelo are available at https://github.com/Tarun-Mahajan/noSpliceVelo

## Supplementary Figures

**Fig. S1:**
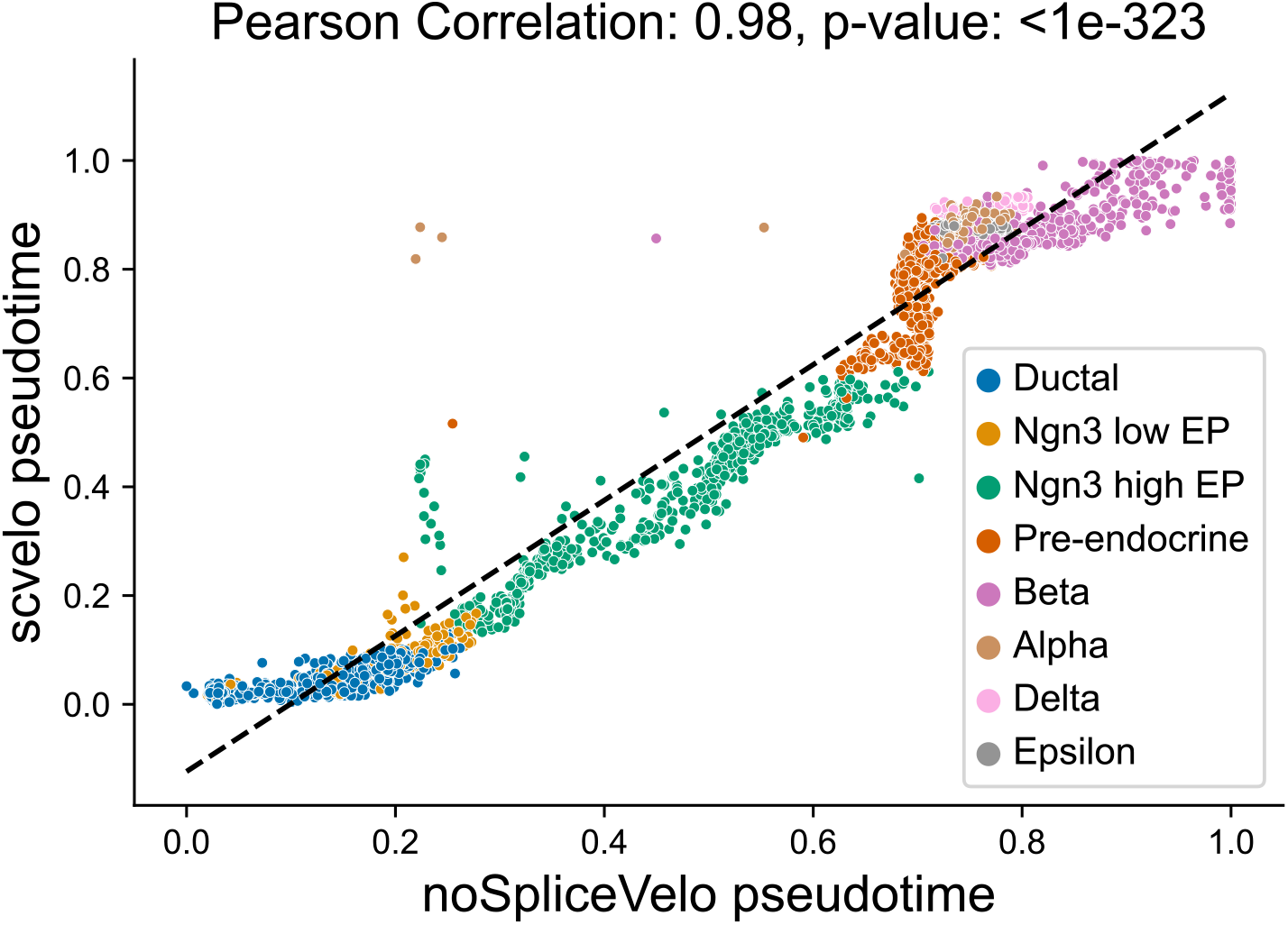
noSpliceVelo’s pseudotime is strongly correlated with scVelo’s pseudotime for mouse pancreatic endocrinogenesis. Scatter plot showing the correlation between noSpliceVelo’s pseudotime (x-axis) and scVelo’s pseudotime (y-axis). The pseudotime values for the *β* and *α* terminal state from scVelo are narrowly distributed around a similar value (flat spread). This shows that scVelo’s pseudotime is unable to assign later pseudotimes to the *β* state.

**Fig. S2:**
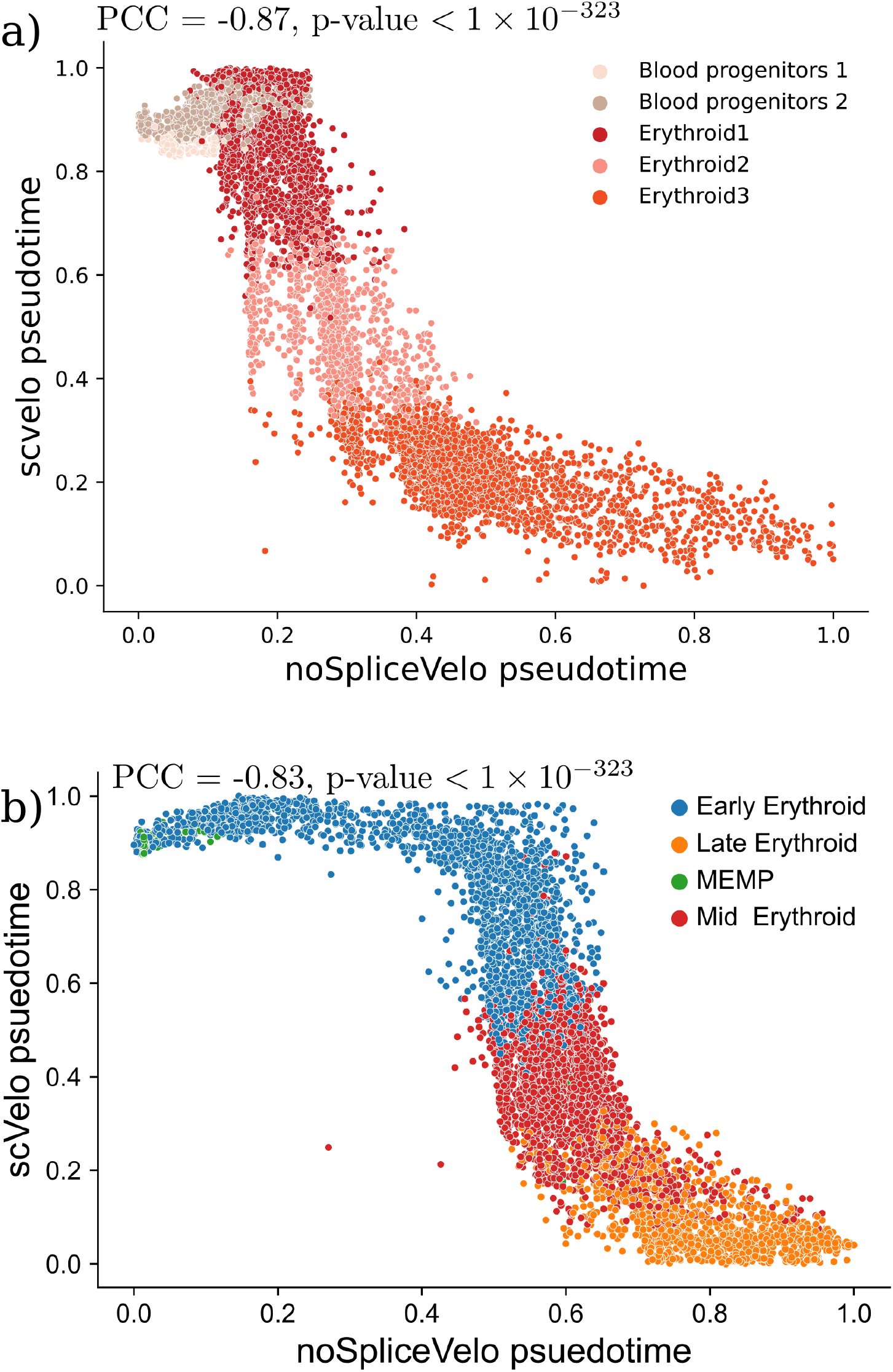
noSpliceVelo’s pseudotime accurately predicts the trajectory, while scVelo’s pseudotime infers biologically implausible flows for mouse and human erythroid maturation. a,b) Scatter plots illustrating the anti-correlation between noSpliceVelo’s pseudotime (x-axis) and scVelo’s pseudotime (y-axis) for a) mouse and b) human erythroid differentiation. noSpliceVelo correctly predicts the flow from progenitor to terminal states through intermediate stages. In contrast, scVelo predicts distorted or reversed trajectories from immature to mature cellular states. The flat regions of the inverse sigmoidal shape indicate that scVelo has poor resolution in pseudotime for the early and late cellular states.

## References

[1] Griffiths, J.A., Scialdone, A., Marioni, J.C.: Using single-cell genomics to understand developmental processes and cell fate decisions. Molecular systems biology 14(4), 8046 (2018)

[2] Kulkarni, A., Anderson, A.G., Merullo, D.P., Konopka, G.: Beyond bulk: a review of single cell transcriptomics methodologies and applications. Current opinion in biotechnology 58, 129–136 (2019)

[3] Tanay, A., Regev, A.: Scaling single-cell genomics from phenomenology to mechanism. Nature 541(7637), 331–338 (2017)

[4] Qiu, Q., Hu, P., Qiu, X., Govek, K.W., Ćamara, P.G., Wu, H.: Massively parallel and time-resolved rna sequencing in single cells with scnt-seq. Nature methods 17(10), 991–1001 (2020)

[5] Cui, H., Maan, H., Vladoiu, M.C., Zhang, J., Taylor, M.D., Wang, B.: Deepvelo: deep learning extends rna velocity to multi-lineage systems with cell-specific kinetics. Genome Biology 25(1), 27 (2024)

[6] Qiu, X., Zhang, Y., Martin-Rufino, J.D., Weng, C., Hosseinzadeh, S., Yang, D., Pogson, A.N., Hein, M.Y., Min, K.H.J., Wang, L., et al.: Mapping transcriptomic vector fields of single cells. Cell 185(4), 690–711 (2022)

[7] Huang, D., Ma, N., Li, X., Gou, Y., Duan, Y., Liu, B., Xia, J., Zhao, X., Wang, X., Li, Q., et al.: Advances in single-cell rna sequencing and its applications in cancer research. Journal of hematology & oncology 16(1), 98 (2023)

[8] Potter, S.S.: Single-cell rna sequencing for the study of development, physiology and disease. Nature Reviews Nephrology 14(8), 479–492 (2018)

[9] Jagadeesh, K.A., Dey, K.K., Montoro, D.T., Mohan, R., Gazal, S., Engreitz, J.M., Xavier, R.J., Price, A.L., Regev, A.: Identifying disease-critical cell types and cellular processes by integrating single-cell rna-sequencing and human genetics. Nature genetics 54(10), 1479–1492 (2022)

[10] Sande, B., Lee, J.S., Mutasa-Gottgens, E., Naughton, B., Bacon, W., Manning, J., Wang, Y., Pollard, J., Mendez, M., Hill, J., et al.: Applications of single-cell rna sequencing in drug discovery and development. Nature Reviews Drug Discovery 22(6), 496–520 (2023)

[11] Heath, J.R., Ribas, A., Mischel, P.S.: Single-cell analysis tools for drug discovery and development. Nature reviews Drug discovery 15(3), 204–216 (2016)

[12] La Manno, G., Soldatov, R., Zeisel, A., Braun, E., Hochgerner, H., Petukhov, V., Lidschreiber, K., Kastriti, M.E., onnerberg, P., Furlan, A., et al.: Rna velocity of single cells. Nature 560(7719), 494–498 (2018)

[13] Bergen, V., Lange, M., Peidli, S., Wolf, F.A., Theis, F.J.: Generalizing rna velocity to transient cell states through dynamical modeling. Nature biotechnology 38(12), 1408–1414 (2020)

[14] Bergen, V., Soldatov, R.A., Kharchenko, P.V., Theis, F.J.: Rna velocity—current challenges and future perspectives. Molecular systems biology 17(8), 10282 (2021)

[15] Qin, Q., Bingham, E., La Manno, G., Langenau, D.M., Pinello, L.: Pyro-velocity: Probabilistic rna velocity inference from single-cell data. bioRxiv, 2022–09 (2022)

[16] Gayoso, A., Weiler, P., Lotfollahi, M., Klein, D., Hong, J., Streets, A., Theis, F.J., Yosef, N.: Deep generative modeling of transcriptional dynamics for rna velocity analysis in single cells. Nature methods 21(1), 50–59 (2024)

[17] Li, S., Zhang, P., Chen, W., Ye, L., Brannan, K.W., Le, N.-T., Abe, J.-i., Cooke, J.P., Wang, G.: A relay velocity model infers cell-dependent rna velocity. Nature biotechnology 42(1), 99–108 (2024)

[18] Gao, M., Qiao, C., Huang, Y.: Unitvelo: temporally unified rna velocity reinforces single-cell trajectory inference. Nature Communications 13(1), 6586 (2022)

[19] Kouadri Boudjelthia, I., Milite, S., El Kazwini, N., Fernandez-Mateos, J., Valeri, N., Huang, Y., Sottoriva, A., Sanguinetti, G.: Neurovelo: interpretable learning of cellular dynamics from single-cell transcriptomic data. bioRxiv, 2023–11 (2023)

[20] Gorin, G., Fang, M., Chari, T., Pachter, L.: Rna velocity unraveled. PLOS Computational Biology 18(9), 1010492 (2022)

[21] Paulsson, J., Berg, O.G., Ehrenberg, M.: Stochastic focusing: fluctuation-enhanced sensitivity of intracellular regulation. Proceedings of the National Academy of Sciences 97(13), 7148–7153 (2000)

[22] Singh, A., Bokes, P.: Consequences of mrna transport on stochastic variability in protein levels. Biophysical journal 103(5), 1087–1096 (2012)

[23] Gorin, G., Pachter, L.: Modeling bursty transcription and splicing with the chemical master equation. Biophysical Journal 121(6), 1056–1069 (2022)

[24] Raj, A., Peskin, C.S., Tranchina, D., Vargas, D.Y., Tyagi, S.: Stochastic mrna synthesis in mammalian cells. PLoS biology 4(10), 309 (2006)

[25] Bastidas-Ponce, A., Tritschler, S., Dony, L., Scheibner, K., Tarquis-Medina, M., Salinno, C., Schirge, S., Burtscher, I., Böttcher, A., Theis, F.J., et al.: Comprehensive single cell mrna profiling reveals a detailed roadmap for pancreatic endocrinogenesis. Development 146(12), 173849 (2019)

[26] Pijuan-Sala, B., Griffiths, J.A., Guibentif, C., Hiscock, T.W., Jawaid, W., Calero-Nieto, F.J., Mulas, C., Ibarra-Soria, X., Tyser, R.C., Ho, D.L.L., et al.: A single-cell molecular map of mouse gastrulation and early organogenesis. Nature 566(7745), 490–495 (2019)

[27] Popescu, D.-M., Botting, R.A., Stephenson, E., Green, K., Webb, S., Jardine, L., Calderbank, E.F., Polanski, K., Goh, I., Efremova, M., et al.: Decoding human fetal liver haematopoiesis. Nature 574(7778), 365–371 (2019)

[28] Rodriguez, J., Larson, D.R.: Transcription in living cells: molecular mechanisms of bursting. Annual review of biochemistry 89(1), 189–212 (2020)

[29] Cao, Z., Grima, R.: Analytical distributions for detailed models of stochastic gene expression in eukaryotic cells. Proceedings of the National Academy of Sciences 117(9), 4682–4692 (2020)

[30] Larsson, A.J., Johnsson, P., Hagemann-Jensen, M., Hartmanis, L., Faridani, O.R., Reinius, B., Segerstolpe, Å., Rivera, C.M., Ren, B., Sandberg, R.: Genomic encoding of transcriptional burst kinetics. Nature 565(7738), 251–254 (2019)

[31] Dar, R.D., Razooky, B.S., Singh, A., Trimeloni, T.V., McCollum, J.M., Cox, C.D., Simpson, M.L., Weinberger, L.S.: Transcriptional burst frequency and burst size are equally modulated across the human genome. Proceedings of the National Academy of Sciences 109(43), 17454–17459 (2012)

[32] Sanchez, A., Golding, I.: Genetic determinants and cellular constraints in noisy gene expression. Science 342(6163), 1188–1193 (2013)

[33] Skinner, S.O., Xu, H., Nagarkar-Jaiswal, S., Freire, P.R., Zwaka, T.P., Golding, I.: Single-cell analysis of transcription kinetics across the cell cycle. Elife 5, 12175 (2016)

[34] McInnes, L., Healy, J., Melville, J.: Umap: Uniform manifold approximation and projection for dimension reduction. arXiv preprint 1802.03426 (2018)

[35] Lopez, R., Regier, J., Cole, M.B., Jordan, M.I., Yosef, N.: Deep generative modeling for single-cell transcriptomics. Nature methods 15(12), 1053–1058 (2018)

[36] Kingma, D.P., Welling, M.: Auto-encoding variational bayes. arXiv preprint 1312.6114 (2013)

[37] Li, J., Pan, X., Yuan, Y., Shen, H.-B.: Tfvelo: gene regulation inspired rna velocity estimation. Nature Communications 15(1), 1387 (2024)

[38] Qiao, C., Huang, Y.: Representation learning of rna velocity reveals robust cell transitions. Proceedings of the National Academy of Sciences 118(49), 2105859118 (2021)

[39] Barile, M., Imaz-Rosshandler, I., Inzani, I., Ghazanfar, S., Nichols, J., Marioni, J.C., Guibentif, C., Göttgens, B.: Coordinated changes in gene expression kinetics underlie both mouse and human erythroid maturation. Genome biology 22, 1–22 (2021)

[40] Ochiai, H., Sugawara, T., Sakuma, T., Yamamoto, T.: Stochastic promoter activation affects nanog expression variability in mouse embryonic stem cells. Scientific reports 4(1), 1–9 (2014)

[41] Ochiai, H., Hayashi, T., Umeda, M., Yoshimura, M., Harada, A., Shimizu, Y., Nakano, K., Saitoh, N., Liu, Z., Yamamoto, T., et al.: Genome-wide kinetic properties of transcriptional bursting in mouse embryonic stem cells. Science advances 6(25), 6699 (2020)

[42] Nicolas, D., Zoller, B., Suter, D.M., Naef, F.: Modulation of transcriptional burst frequency by histone acetylation. Proceedings of the National Academy of Sciences 115(27), 7153–7158 (2018)

[43] Meeussen, J.V., Lenstra, T.L.: Time will tell: comparing timescales to gain insight into transcriptional bursting. Trends in Genetics 40(2), 160–174 (2024)

[44] Beckman, W.F., Jiménez, M.Á.L., Verschure, P.J.: Transcription bursting and epigenetic plasticity: an updated view. Epigenetics Communications 1(1), 6 (2021)

[45] Brouwer, I., Kerklingh, E., Leeuwen, F., Lenstra, T.L.: Dynamic epistasis analysis reveals how chromatin remodeling regulates transcriptional bursting. Nature structural & molecular biology 30(5), 692–702 (2023)

[46] Leyes Porello, E.A., Trudeau, R.T., Lim, B.: Transcriptional bursting: stochasticity in deterministic development. Development 150(12), 201546 (2023)

[47] Tedesco, M., Giannese, F., Lazarević, D., Giansanti, V., Rosano, D., Monzani, S., Catalano, I., Grassi, E., Zanella, E.R., Botrugno, O.A., et al.: Chromatin velocity reveals epigenetic dynamics by single-cell profiling of heterochromatin and euchromatin. Nature biotechnology 40(2), 235–244 (2022)

[48] Felce, C., Gorin, G., Pachter, L.: A biophysical model for atac-seq data analysis. bioRxiv, 2024–01 (2024)

[49] Sarkar, A., Stephens, M.: Separating measurement and expression models clarifies confusion in single-cell rna sequencing analysis. Nature genetics 53(6), 770–777 (2021)

[50] Tang, W., Bertaux, F., Thomas, P., Stefanelli, C., Saint, M., Marguerat, S., Shahrezaei, V.: baynorm: Bayesian gene expression recovery, imputation and normalization for single-cell rna-sequencing data. Bioinformatics 36(4), 1174–1181 (2020)

[51] Tang, W., Jørgensen, A.C.S., Marguerat, S., Thomas, P., Shahrezaei, V.: Modelling capture efficiency of single-cell rna-sequencing data improves inference of transcriptome-wide burst kinetics. Bioinformatics 39(7), 395 (2023)

[52] Svensson, V.: Droplet scrna-seq is not zero-inflated. Nature Biotechnology 38(2), 147–150 (2020)

[53] Li, T., Shi, J., Wu, Y., Zhou, P.: On the mathematics of rna velocity i: Theoretical analysis. CSIAM Transactions on Applied Mathematics 2 (2021) 10.4208/csiam-am.SO-2020-0001

[54] Blei, D.M., Kucukelbir, A., McAuliffe, J.D.: Variational inference: A review for statisticians. Journal of the American statistical Association 112(518), 859–877 (2017)

[55] Wolf, F.A., Angerer, P., Theis, F.J.: Scanpy: large-scale single-cell gene expression data analysis. Genome biology 19, 1–5 (2018)

